# Inhibition of MMP 2&9 protects the endothelial glycocalyx and improves diastolic function in diabetic cardiomyopathy

**DOI:** 10.1101/2023.12.18.572183

**Authors:** Stanley Buffonge, Yan Qiu, Sarah Fawaz, Monica Gamez, Michael Crompton, Matthew J Butler, Gavin I Welsh, Paolo Madeddu, Rebecca R Foster, Simon C Satchell

## Abstract

The coronary microvascular endothelial glycocalyx (EGlx) is a vital regulator of vascular permeability and EGlx damage contributes to the development of diabetic cardiomyopathy. Matrix metalloproteinases 2 and 9 (MMP2/9) have been identified as key enzymes in the degradation of EGlx components, notably syndecan 4 (SDC4), and are upregulated in diabetes. We tested the hypothesis that inhibition of MMP2/9 can protect the EGlx and improve diastolic function in diabetic cardiomyopathy. Type 1 diabetes was induced in FVB mice by streptozotocin (STZ) injections. Mice were treated with daily injections of the MMP2/9 inhibitor, SB-3CT, for 2 weeks from 7 weeks post STZ. Echocardiography was utilised to assess heart function and lectin staining for the measurement of EGlx depth. Immunolabelling of heart sections for albumin provided an indication of albumin extravasation. A mechanism of EGlx shedding was investigated in vitro in human coronary microvascular endothelial cells treated with TNF-α and SB-3CT. Diabetic mice developed diastolic dysfunction from 6 weeks post STZ. MMP2/9 inhibition reversed diastolic dysfunction, EGlx thinning and albumin extravasation in diabetic animals. In vitro, TNF-α caused an increase in MMP9 activity and SDC4 shedding from human coronary microvascular endothelial cells. Treatment with SB-3CT reduced MMP9 activity and prevented SDC4 shedding. Knockdown of MMP9 expression prevented TNF-α induced SDC4 shedding. This study demonstrates MMP2/9 inhibition as a strategy to protect the EGlx and improve diastolic function in diabetic cardiomyopathy. Our findings suggest new avenues for therapeutic interventions in cardiovascular complications associated with diabetes.

**Statements and Declarations:** *Competing interests:* The authors have no competing interests to declare that are relevant to the content of this article.

## Introduction

In 1974, the Framingham study revealed that the likelihood of the development of heart failure was increased in those with diabetes than in those without [15]. Today it is understood that diabetes is an independent risk factor for heart failure and the term diabetic cardiomyopathy (DCM) has now been coined. In fact, DCM was first described in 1972 as a result of post-mortem findings from diabetic patients who had heart failure symptoms, in the absence of any other cardiovascular diseases [34]. DCM is clinically characterised by left ventricular diastolic dysfunction [23] in the absence of systolic dysfunction. Ultimately, DCM is the progression of diastolic dysfunction to heart failure, independently of other cardiovascular factors such as hypertension and coronary artery disease, in patients with diabetes [22]. With a fivefold increase in the risk of heart failure in diabetic patients, DCM is now recognised as a major cardiovascular complication in patients with diabetes [8]. The mechanisms of DCM are complex and point to various potential therapeutic targets. Despite this, there is currently a lack of treatments specifically for DCM.

Microvascular damage is an early characteristic of DCM, and this is associated with altered permeability. The heart is extremely sensitive to changes in microvascular permeability, with even a 2% increase in interstitial fluid compromising heart function [9]. In fact, increased microvascular permeability, independently of other factors such as inflammation and fibrosis, has been shown to cause diastolic dysfunction [1]. A vital regulator of microvascular permeability is the endothelial glycocalyx (EGlx). This is a brush-like layer that lines the luminal side of the vascular endothelial cells [19], thus providing an interface between the flowing blood and the endothelial cell membranes. This heterogeneous layer is largely composed of proteoglycans and glycoproteins, which adsorb additional plasma components. Proteoglycans consist of a cell-surface anchored core protein with one or more negatively charged glycosaminoglycan (GAG) side chains [42]. Syndecans are transmembrane core proteins [33] and are the most abundant core proteins of the proteoglycan family within the EGlx with the largest attachment possibilities for GAGs when compared to the other core proteins [27].

It is known that diabetes causes damage to the EGlx [5, 20, 28–30, 38] and we have recently shown coronary microvascular EGlx damage, for the first time, in mice with DCM [30]. In our recent study we have shown that preservation of the coronary microvascular EGlx with angiopoietin 1 reduced perivascular oedema and improved diastolic function in both type 1 and 2 models of DCM [30]. Therefore, we have shown that protecting the EGlx is crucial in DCM. An important contributor to EGlx damage is matrix metalloproteinases (MMPs) [31, 32]. Previously, it has been shown that diabetes results in an upregulation of MMP9 activity resulting in damage to the glomerular EGlx [32]. As well as this an increase in MMP9 activity has been identified in models of DCM [7, 30]. It has also been shown that tumour necrosis factor a (TNF-α), a proinflammatory cytokine which is upregulated in diabetes, leads to increased MMP9 activity and causes shedding of syndecan 4 (SDC4) [31].

Recognizing the pivotal role of the EGlx in vascular permeability and its impact on heart function, our hypothesis posits that DCM induces an upregulation in MMP9 activity, resulting in damage to the coronary microvascular EGlx and subsequent diastolic dysfunction. Consequently, we propose that inhibition of MMP2/9 will serve as a protective measure for the EGlx, ultimately improving diastolic function. This hypothesis not only sheds light on the intricate interplay between diabetes, MMPs, and the EGlx but also opens avenues for targeted therapeutic interventions in the realm of DCM.

## Methods

### Animal model

Animal studies were conducted in accordance with local ethical standards. All animal procedures performed conformed to the guidelines from Directive 2010/63/EU of the European Parliament on the protection of animals used for scientific purposes and were approved by the University of Bristol and the British Home Office (PPL: P855B71B4).

Type 1 diabetes was induced in 6-week-old male FVB mice (FVB/NCrl, Charles River) by injection of streptozotocin (STZ) (S0130-500MG; Sigma Aldrich) at 50 mg/kg body weight per day, i.p. for 5 consecutive days. All animals were fasted for 4–6h before the administration of STZ. Control animals received citric acid buffer or were not injected. Hyperglycaemia was confirmed by blood glucose levels of more than 16 mmol/L on two consecutive days. Body weight was monitored weekly and blood glucose was assessed with a glucometer (ACCU-CHEK Aviva, Roche, UK) at 2, 6, and 9 weeks post the induction of diabetes.

### MMP inhibitor treatment

From 7 weeks post the induction of diabetes, mice were treated with the MMP2 and 9 specific inhibitor [11, 14, 41, 46], SB-3CT (MedChemExpress, HY-12354) at 10mg/kg in 10% DMSO, 40% PEG-300, 5% tween-80 and 45% saline, daily for 2 weeks. The dosage was chosen based on previous literature which utilised SB-3CT in mice [3, 41]. Diabetic mice were randomized to receive either SB-3CT or the vehicle leading to three final groups: control, diabetes, and diabetes + SB-3CT. All mice in the control and diabetes groups received the vehicle alone daily.

### Haemodynamic measurements

Echocardiography (Vevo 3100, Visual Sonics, Toronto, Canada) was utilized to assess diastolic and systolic function as described previously [30]. In brief, diastolic function was determined with pulsed wave Doppler and tissue Doppler to measure the E/A ratio, isovolumetric relaxation time (IVRT) and E’/A’. Images were captured at a heart rate of 380±20 beats per min under 1-3% isoflurane anaesthesia. Systolic function was determined with the analysis of M-mode images at a heart rate of 450±30 beats per minute. All measurements were captured within 30 minutes of the mouse being under anaesthesia.

### Tissue collection

At the end of the study, whilst under 1-3% isoflurane mice were perfused through the apex of the left ventricle with cadmium chloride, to stop the heart at diastole, followed by PBS as described previously [30]. Anaesthesia was monitored carefully by observing the respiratory rate and heart rate as well as by pedal reflex (firm pinch toe). When the heart expanded as a result of cadmium chloride, the right atrium was snipped, and blood was collected in 1.5 ml Eppendorf tubes coated with heparin (764 USP units/ml). The heart was taken and either PFA fixed or frozen in liquid nitrogen.

### Staining the EGlx by lectin-based immunofluorescence

To determine the EGlx depth, the tissue sections (5μm) were first dewaxed in xylene for 20min followed by rehydration of the tissue in graded ethanol (100%, 90%, and 50%). The sections were then washed 3 times for 5min with PBS before being blocked with blocking solution (1% BSA in PBS containing 0.5% tween-20) for 1h at room temperature, followed by washes 3 times with PBX (0.5% tween in PBS). For endogenous biotin blocking, the sections were incubated with streptavidin (ZJ0912, BioServ UK Limited) for 15min at room temperature. Following this, the sections were washed once with PBX and then incubated with biotin (ZJ0912; BioServ UK Limited) for 15min at room temperature. After another set of washes with PBX (2x), the sections were incubated with biotinylated Sambucus Nigra Lectin (SNA) in 1% BSA/PBX at pH 6.8 (ZF0207; Vector Laboratories) for 1h at room temperature. This was followed by 4 washes in PBX before being incubated with the secondary antibody streptavidin-488 (2273715, Thermo Fisher Scientific; 1% BSA/0.5% PBX; Ph 6.8) for 1h at room temperature. After 2 more washes with PBX followed by 2 washes with PBS, the sections were incubated with 4’,6-diamidino-2-phenylindole (DAPI; D1306, Thermo Fisher Scientific; 1:1000 in PBS) for 2min at room temperature to counterstain the nucleus before being washed twice in PBS. To stain the membrane, sections were incubated with Octadecyl Rhodamine B Chloride (R18; 2061441, Invitrogen; 1:1000 in PBS) for 10min at room temperature. No washes occurred after this incubation, but the sections were briefly rinsed with PBS. Images were acquired by confocal microscopy (Confocal 13, Leica SP8) at 63x magnification at the Wolfson Bioimaging Facility at the University of Bristol.

### Quantification of EGlx depth

Analysis of EGlx depth was performed using a macro in Image J which measures the anatomical distance between the fluorescence intensity peaks of lectin and R18 as an indication of EGlx depth as described previously [5]. Capillary (3-10 μm in diameter) cross-sections were selected for analysis by circularity (ratio of shortest and longest diameter, 0.8 to 1). A perpendicular profile line was drawn from the inside to the outside of a capillary loop crossing the lectin labelled EGlx first, followed by the R18 labelled endothelial membrane. Fluorescence intensity plot profiles were then generated for the lectin-labelled components of the EGlx and endothelial cell membrane label. The distance between the peak signals from the lectin-488 and the R18 labels (peak-to-peak) was used as an indication of EGlx thickness. This method was automated using an Image J (NIH) macro to take multiple measurements in a preselected capillary loop as described previously [5]. Gaussian curves were applied to the raw intensity data of each plot for peak-to-peak measurements. The mean was subsequently determined from 360 lines per capillary cross section, and 15 capillaries per heart. Data were excluded with a standard deviation of more than 7.5 and/or a signal-to-noise ratio of less than 15 from the 360 measurements of the capillary loops.

### Albumin Immunofluorescence

Heart sections were deparaffinised and rehydrated in graded ethanol. The tissue was blocked with 0.3% Triton-X, 3% BSA in PBS solution for 30min and then incubated with goat anti-mouse albumin antibody (A90-134A-17; Bethyl Laboratories) at 1:200 in blocking buffer overnight at 4°C in a humidity chamber. Sections were washed three times for 5min each in 0.3% Triton-X in PBS. Tissue sections were then incubated with donkey anti-goat Alexa Fluor-488 (15930877; Fisher Scientific Ltd) at 1:200 in blocking buffer for 1h at room temperature. Images were captured with a confocal microscope (Confocal 13, Leica SP8) and the corrected total fluorescence was calculated using the following equation:

CTCF = Integrated density – (area of the region of interest x mean fluorescence of background means)

### Cell culture

Human conditionally immortalised coronary microvascular endothelial cells (ciCMVEC) were established as described previously [30], using a similar approach as that described for glomerular endothelial cells [35]. In brief, primary CMVEC were transduced with temperature-sensitive simian virus 40 large tumor (tsSV40LT) antigen and telomerase using retroviral vectors allowing the proliferation of the cells at 33°C (without telomere shortening). Inactivation of the virus at 37°C, allows the cells to become quiescent. Cells were thermo-switched to 37°C once they were at 80% confluency for 3-5 days to allow transgene inactivation for experimental use. ciCMVEC were cultured in endothelial growth medium 2 microvascular (EGM2-MV; Lonza, Basel, Switzerland) containing 5% fetal calf serum but without vascular endothelial growth factor A or Gentamicin Sulfate-Amphotericin.

### Treatment of ciCMVEC with TNF-α and SB-3CT

To investigate the effects of TNF-α on the EGlx as well its effect on MMP regulation, ciCMVEC were placed in serum-free media 1h before treatment with 10 ng/ml of Recombinant Human TNF-α Protein (210-TA-005; R&D systems) or vehicle (PBS) for 6h. To investigate the effects of MMP inhibition on the EGlx, some cells were treated with 10mM of the MMP2 and 9 inhibitor, SB-3CT (HY-12354; MedChemExpress) whilst the others were treated with the vehicle (DMSO) 2h prior to TNF-α treatment and remaining for the duration of TNF-α treatment.

### Knockdown of MMP9 expression in ciCMVEC

To specifically investigate the role of MMP9 on EGlx shedding, MMP9 expression was knocked down in ciCMVEC using MMP9 shRNA lentiviral particles (sc-29400-V, Santa Cruz Biotechnology). Controls were cells exposed to lentiviral particles with a scrambled shRNA sequence. Cells were seeded in 6 well plates at a density of 31,250 cells/cm^2^. Transfected cells were treated with TNF-α as described above for 6h.

### RNA extraction, cDNA conversion and Quantitative polymerase chain reaction (qPCR)

Monolayers of ciCMVEC grown in 6 well plates were lysed, and RNA was extracted using a RNeasy mini kit (74104, Qiagen) according to the manufacturer’s protocol. One μg RNA was converted to cDNA using a highcapacity RNA to cDNA kit (4387406, Applied Biosystems) according to the manufacturer’s instructions.

mRNA expression was quantified using a StepOnePlus Real-Time PCR system (Applied Biosystem). The master mix was made using 61% of SYBR Green (4385612; Thermo Fisher Scientific), 8.9% of the primer mix and 30% of DEPC water. The sample’s CT values were normalised to glyceraldehyde-3-phosphate dehydrogenase (GAPDH), and further normalised to the control condition. The 2^-ΔΔCT^ method of quantification was used to calculate the fold change of the gene of interest. Primers used in this study are shown in Table 1.

**Table 1.**
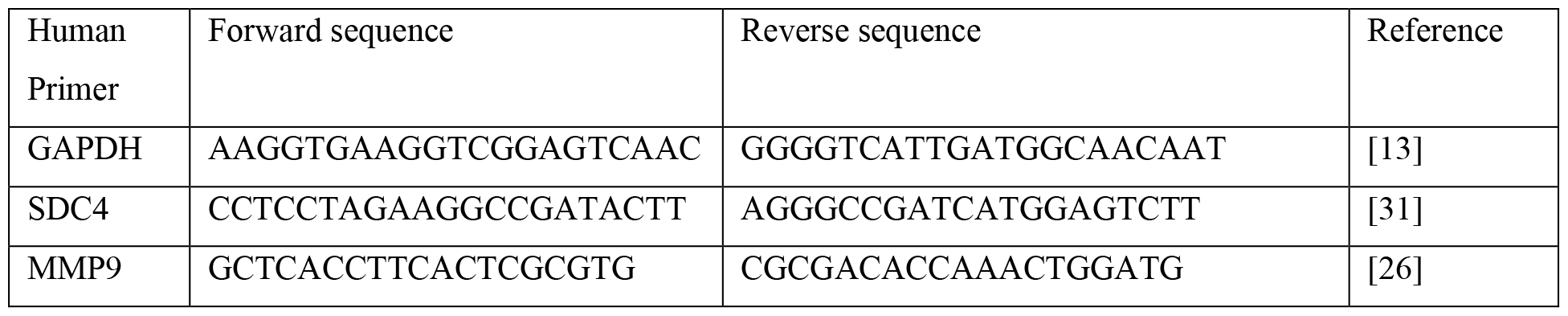
Primer sequences. All primers were synthesised by Eurofins genomics. Primers were prepared to a working stock concentration of 10μM.

### SDC4 ELISA

SDC4, shed from ciCMVEC, was quantified in cell culture media using a SDC4 ELISA as per the manufacturer’s instructions (DY2918, R&D Systems).

### Immunofluorescence of SDC4

ciCMVEC were grown on glass coverslips and treated with TNF-α and SB-3CT, as described above. After treatment, the cells were fixed for 10min with 4% PFA and blocked with 1% BSA in PBS for 1h. The cells were incubated with an anti-SDC4 primary antibody at 10 mg/ml in blocking buffer (AF2918-SP, Bio-Techne) for 1h at room temperature. After three washes with PBS for 5min each, the cells were incubated with the Anti Goat IgG/Alexa Fluor 488 secondary antibody (Fisher Scientific Ltd, 15930877; 1:500 in blocking buffer) for 1h at room temperature and the washes were repeated. To label the nucleus, the cells were incubated with DAPI, (1:1000 in PBS) for 2min at room temperature before being washed three times with PBS. The cells were mounted with vectashield antifade mounting medium (BioServ UK Limited) inverted onto a microscope slide. Three images were taken at random areas for each sample using a confocal microscope. A mean fluorescent intensity for each sample was calculated from the total fluorescent intensity from each image and normalised to DAPI to correct for cell number using Image J [36].

### Shed sulphated GAG Assay

An Alcian blue colorimetric assay was used to quantify sulphated GAG shed into the culture media from the ciCMVEC surface [31]. After treatment with TNF-α, as described above, the media was harvested and centrifuged at 800g for 3min to remove cellular debris. The supernatant was added to a freshly prepared solution of Alcian blue (0.4%) in sodium acetate (0.5M) (33864-99-2; Sigma-Aldrich), magnesium chloride hexahydrate (30 mM), and sulphuric acid (2.8%). Standards were prepared using chondroitin sulfate (CS, C4384; Sigma-Aldrich) and absorbance at 490nm was measured using a plate reader after 15min of incubation. The linear relation between GAG mass and decreased absorption of 490nm by Alcian blue solution was used to quantify supernatant GAG content, referenced to known concentrations (0 to 125μg/ml) of CS.

### Quantification of MMP activity

MMP2 and 9 activity assays were used to measure MMP2 and 9 activities in frozen mouse heart tissue and plasma following the manufacturer’s instructions (AS-72017, Eurogentec). MMP9 activity in conditioned media from human ciCMVEC was also assessed.

### Statistical Analysis

All statistical analyses were performed with Prism version 9 (GraphPad Software) (^*^p<0.05, ^**^p<0.01, ^***^p<0.001, ^****^p<0.0001). Data was first assessed for normal distribution using the Shapiro-Wilk test. For normally distributed data with two groups, statistical significance was determined using an unpaired t-test. Non-normally distributed data was assessed using a Mann-Whitney test. For data with more than two groups, normally distributed data were assessed using ANOVA followed by Tukey’s to compare the means of each group with that of every other group. Non-normally distributed data was assessed using the Kruskal-Wallis test followed by Dunn’s multiple comparison test. To determine if there was any significant correlation between data sets, the Pearson r test was conducted.

## Results

### Mice develop diastolic dysfunction from 6 weeks post the induction of diabetes

Mice treated with STZ developed hyperglycaemia indicated by blood glucose levels higher than 16 mmol/L 2 weeks post the induction of diabetes (Fig. 1a). Diabetic mice gained significantly less weight than control mice throughout the duration of the study (Fig. 1b). Echocardiography was used to assess systolic and diastolic function. As the E/A ratio is consistently reduced in type 1 mouse models of diabetes [6, 30], we assessed this as a primary measure of diastolic function. Diastolic dysfunction was observed in diabetic mice 6 weeks post the induction of diabetes as indicated by a reduced E/A ratio (Fig. 1e). An increased isovolumetric relaxation time (IVRT) was also found providing further evidence of diastolic dysfunction (Fig. 1f) as well as a reduced E’/A’ (Fig. 1g). No systolic dysfunction was detected in diabetic mice (Table 2).

**Table 2:** Mice do not develop systolic dysfunction 6 weeks after induction of diabetes. Systolic function was assessed using M-mode. No significant difference in any systolic parameters was recorded between the controls and diabetic mice (n=7 for control and 9 for diabetes for all parameters; Ejection Fraction: p=0.091; Fractional shortening: p=0.098; Cardiac output: p=0.86)

| Mouse | 6 weeks post diabetes induction |  |  |
| --- | --- | --- | --- |
|  | Ejection Fraction | Fractional Shortening | Cardiac Output |
| Control | 62.99 ± 2.06 | 33.63 ± 1.52 | 17.94 ± 1.03 |
| Diabetes | 69.51 ± 2.72 | 38.83 ± 2.28 | 17.67 ± 1.06 |

**Fig. 1.**
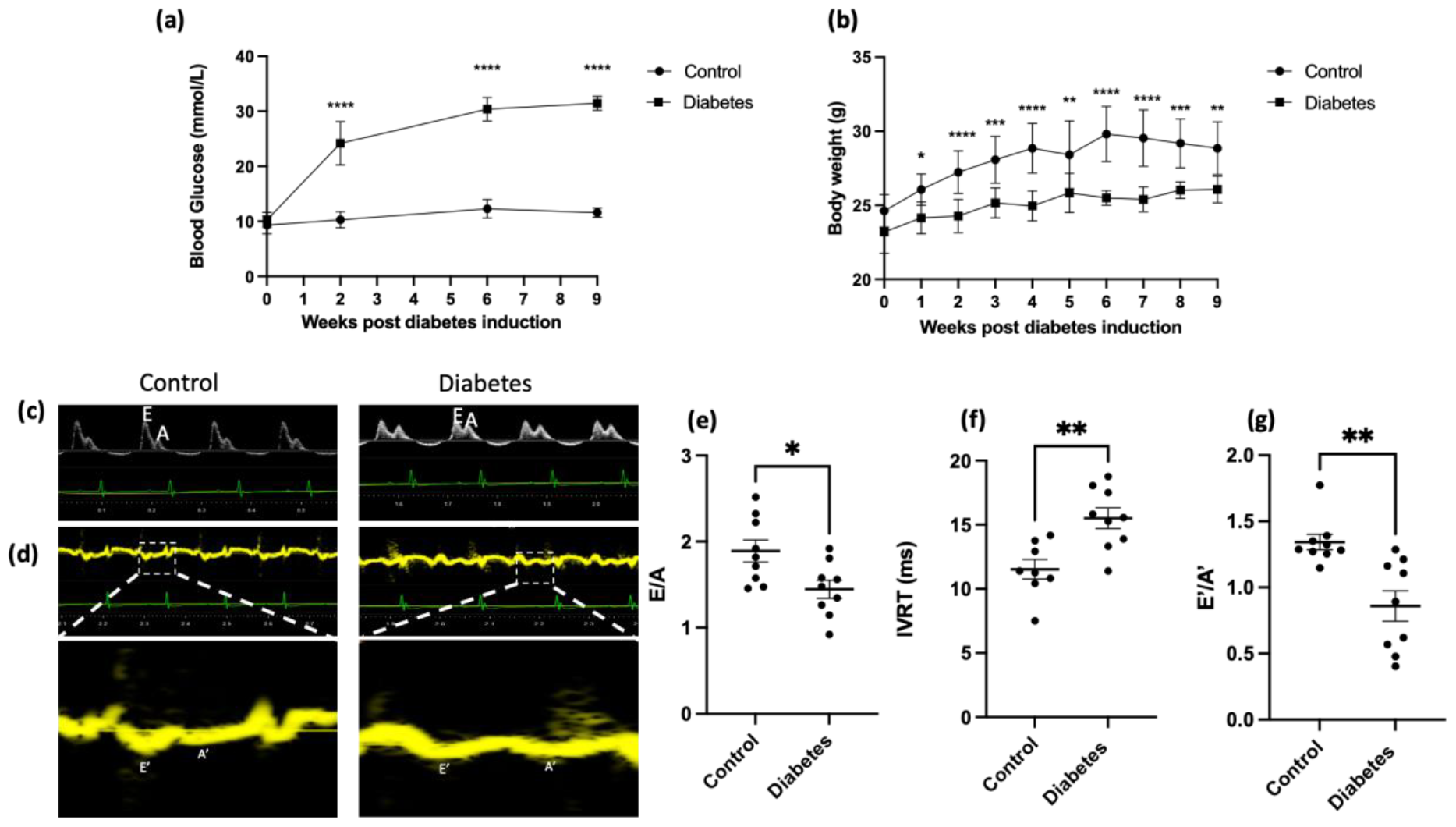
Mice develop diastolic dysfunction 6 weeks post induction of diabetes. (a) Blood glucose post diabetes induction (2 and 6 weeks post diabetes induction: n=10 for control and diabetes; ****p<0.0001; unpaired t test, 9 weeks post diabetes induction: n=10 for control and 5 for diabetes; ****p<0.0001; unpaired t test). (b) Weekly body weight of mice (n=10 for controls for all weeks, 10 for diabetes group from baseline to 6 weeks post diabetes induction, and 5 for diabetes group from 7-9 weeks post diabetes induction; 1 week post diabetes induction: *p<0.05; 2, 4, 6, and 7 weeks post diabetes induction: ****p < 0.0001; 3 and 8 weeks post diabetes induction: ***p < 0.001; 6 and 9 weeks post diabetes induction: **p < 0.01; Twoway ANOVA). (c) Representative pulsed wave Doppler images. (d) Representative tissue Doppler images. (e) Reduced E/A ratio in diabetic mice (n=9 for control and 9 for diabetes; *p<0.05; unpaired t test). (f) Increased IVRT in diabetic mice (n=9 for control and 9 for diabetes; **p<0.01; unpaired t test). (g) Reduced E’/A’ in diabetic mice (n=9 for control and 9 for diabetes; **p<0.01; Mann-Whitney test)

### MMP2/9 inhibition improves diastolic function in diabetic FVB mice

Once diastolic dysfunction was developed in diabetic mice, we investigated if the inhibition of MMP2/9 improves diastolic function in diabetic mice. From 7 weeks post the induction of diabetes, the mice were treated with SB-3CT daily for 2 weeks. Echocardiography was carried out at 9 weeks post diabetes induction to assess systolic and diastolic function. As expected, diastolic dysfunction, indicated by a reduced E/A ratio, increased isovolumetric relaxation time (IVRT) and reduced E’/A’, was observed in diabetic mice. When treated with SB-3CT, diastolic function was improved as shown by an increased E/A ratio, reduced IVRT and increased E’/A’(Fig. 2). SB-3CT had no significant impact on blood glucose with mice remaining hyperglycaemic after MMP inhibition (Figure 2e). No significant difference in systolic function was found between mice 9 weeks post the induction of diabetes (Table 3).

**Table 3:** Mice do not develop systolic dysfunction 9 weeks after induction of diabetes. Systolic function was assessed using M-mode. No significant difference in any systolic parameters was recorded between the controls and diabetic mice (control: n=9 for control, 5 for diabetes, and 5 for diabetes + SB-3CT; Ejection fraction: control vs diabetes, p=0.98, diabetes vs diabetes + SB-3CT, p=0.58; Fractional shortening: control vs diabetes p=0.83, diabetes vs diabetes + SB-3CT, p=0.58; Cardiac output: control vs diabetes, p=0.93; diabetes vs diabetes + SB-3CT, p=0.4; One-way ANOVA)

| Mouse | 9 weeks post diabetes induction |  |  |
| --- | --- | --- | --- |
|  | Ejection Fraction | Fractional Shortening | Cardiac Output |
| Control | 66.36 ± 1.91 | 35.03 ± 1.78 | 17.57 ± 0.85 |
| Diabetes | 66.92 ± 2.55 | 36.51 ± 1.98 | 16.99 ± 1.47 |
| Diabetes + SB-3CT | 63.39 ± 1.6 | 33.59 ± 1.22 | 14.47 ± 1.52 |

**Fig. 2.**
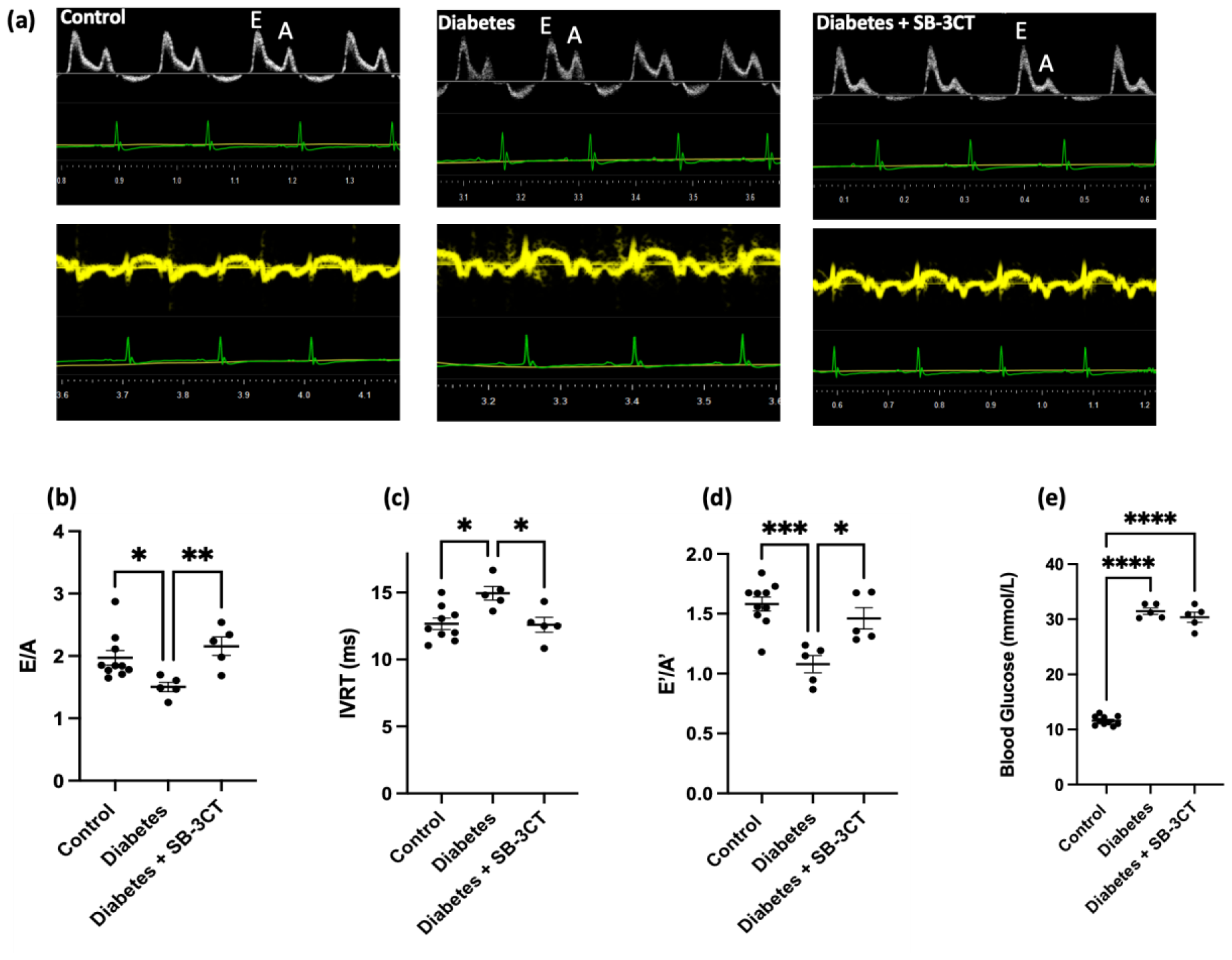
MMP2/9 inhibition improves diastolic function in diabetic FVB mice. Echocardiography was used to assess diastolic function through pulsed wave Doppler and tissue Doppler. (a) Representative pulse wave and tissue Doppler images. (b) E/A ratio determined by pulsed wave Doppler (n=10 for control, 5 for diabetes and 5 for diabetes + SB-3CT; control vs diabetes, *p<0.05; diabetes vs diabetes + SB-3CT, **p<0.01; Kruskal-Wallis test). (c) IVRT determined from pulsed wave Doppler (n=9 for control, 5 for diabetes and 5 for diabetes + SB-3CT; control vs diabetes, *p<0.05; diabetes vs diabetes + SB-3CT, *p<0.05; One-way ANOVA). (d) E’/A’ ratio determined by tissue Doppler (n=10 for control, 5 for diabetes and 5 for diabetes + SB-3CT; Control vs diabetes, ***p<0.001; diabetes vs diabetes + SB-3CT, *p<0.05; One-way ANOVA). (e) Blood Glucose determined from the tail vain (n=10 for control, 5 for diabetes and 5 for diabetes + SB-3CT; control vs diabetes and control vs diabetes + SB-3CT, ****p<0.0001; One-way ANOVA)

### MMP2/9 inhibition protects the coronary microvascular EGlx in diabetic mouse hearts

We next aimed to investigate if the coronary microvascular EGlx depth was reduced in diabetes and if MMP2/9 inhibition protected the EGlx. A measure of EGlx depth was provided using the fluorescence profile peak-to-peak analysis as previously used ex vivo [2, 5, 32] (Fig. 3a.ii). EGlx depth was reduced in diabetic mice 9 weeks after the induction of diabetes and restored when treated with SB-3CT (Fig. 3b). A positive correlation was also found between EGlx depth and E/A ratio (Fig. 3c) indicating that a thicker EGlx is correlated with better diastolic function.

**Fig. 3.**
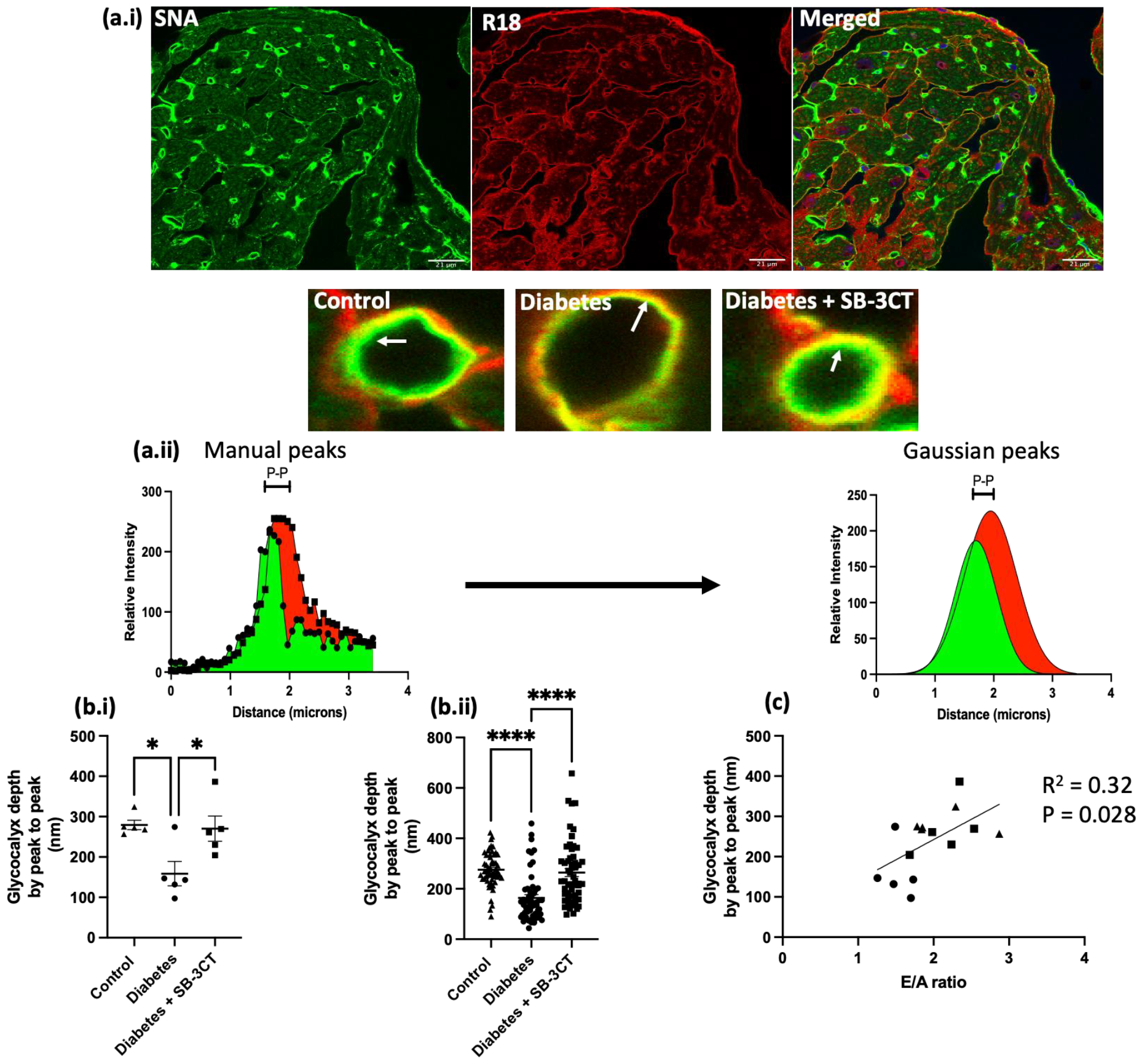
MMP2/9 inhibition protects the coronary microvascular EGlx in diabetic mouse hearts. Paraffin-embedded tissue was sectioned and stained for the EGlx using SNA lectin. (a.i) Representative images of heart tissue stained with SNA lectin (green) and the membrane stained with R18 (red) (scale bar is 21 μm). The white arrow points to the EGlx on the luminal side of the vessel. A line was drawn across the vessel from the centre to generate fluorescent plot profiles and the distance between the SNA peak and the R18 peak was used as an indication of EGlx depth (a.ii). (b.i) Coronary microvascular EGlx depth (n=5 for all groups; *p<0.05; one-way ANOVA). (b.ii) All the vessels that were analysed for EGlx depth (****p<0.0001). (C) Correlation of coronary microvascular EGlx depth with E/A ratio (n=15; R^2^ =0.32; *p<0.05; Pearson r). Triangles represent controls, circles represent diabetes, and squares represent diabetes + SB-3CT

### Increased albumin extravasation in diabetic mouse hearts is ameliorated by MMP inhibition

It is expected that a reduction in EGlx integrity will result in an alteration in microvascular permeability. We have previously shown in human CMVEC that the EGlx is vital for endothelial barrier properties and when we strip the EGlx with enzyme treatment, there is a significant increase in albumin leak across the cell monolayer [30]. To extend this to an in vivo model of DCM and provide an indication of coronary microvascular permeability, mouse heart tissue sections were stained for extravasated albumin. Images of the left ventricle were captured, and fluorescence intensity was quantified. In diabetic mice, there was an increase in interstitial albumin, indicating albumin extravasation due to increased microvascular permeability (Fig. 4aii). This was reduced by SB-3CT treatment. A negative correlation between the albumin intensity and E/A ratio (Fig. 4b) as well as between albumin intensity and EGlx depth (Fig. 4c) was also observed. It can therefore be interpreted that reduced EGlx depth leads to increased microvascular permeability and thus increased albumin extravasation resulting in diastolic dysfunction.

**Fig. 4.**
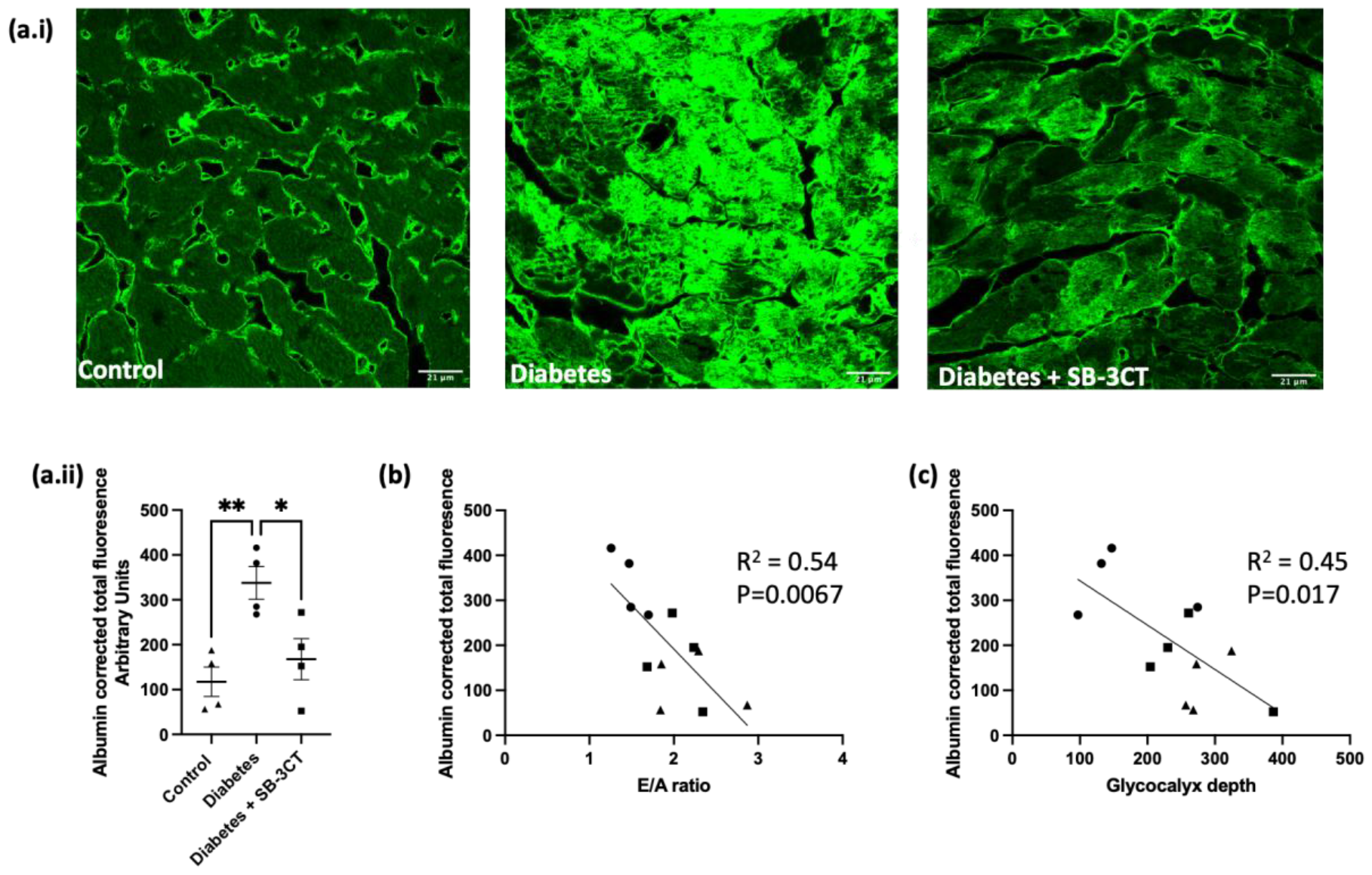
Increased albumin extravasation in diabetic mouse hearts is reduced with SB-3CT. Immunolabelling for albumin in control, diabetes, and diabetes + SB-3CT mouse heart sections. (a.i) Representative images of albumin staining in control, diabetes, and diabetes + SB-3CT (scale bar is 21 μm). (a.ii) Total fluorescence of albumin corrected to the area with background removed (n=4 for all groups; control vs diabetes, **p<0.01; diabetes vs diabetes + SB-3CT, *p<0.05; One-way ANOVA). (b) Negative correlation between albumin fluorescent intensity and E/A ratio (n=12; R^2^=0.54; p**<0.01; Pearson r). (c) Negative correlation between albumin fluorescent intensity and EGlx depth (n=12; R^2^=0.45; p*<0.05; Pearson r)

**Fig. 5.**
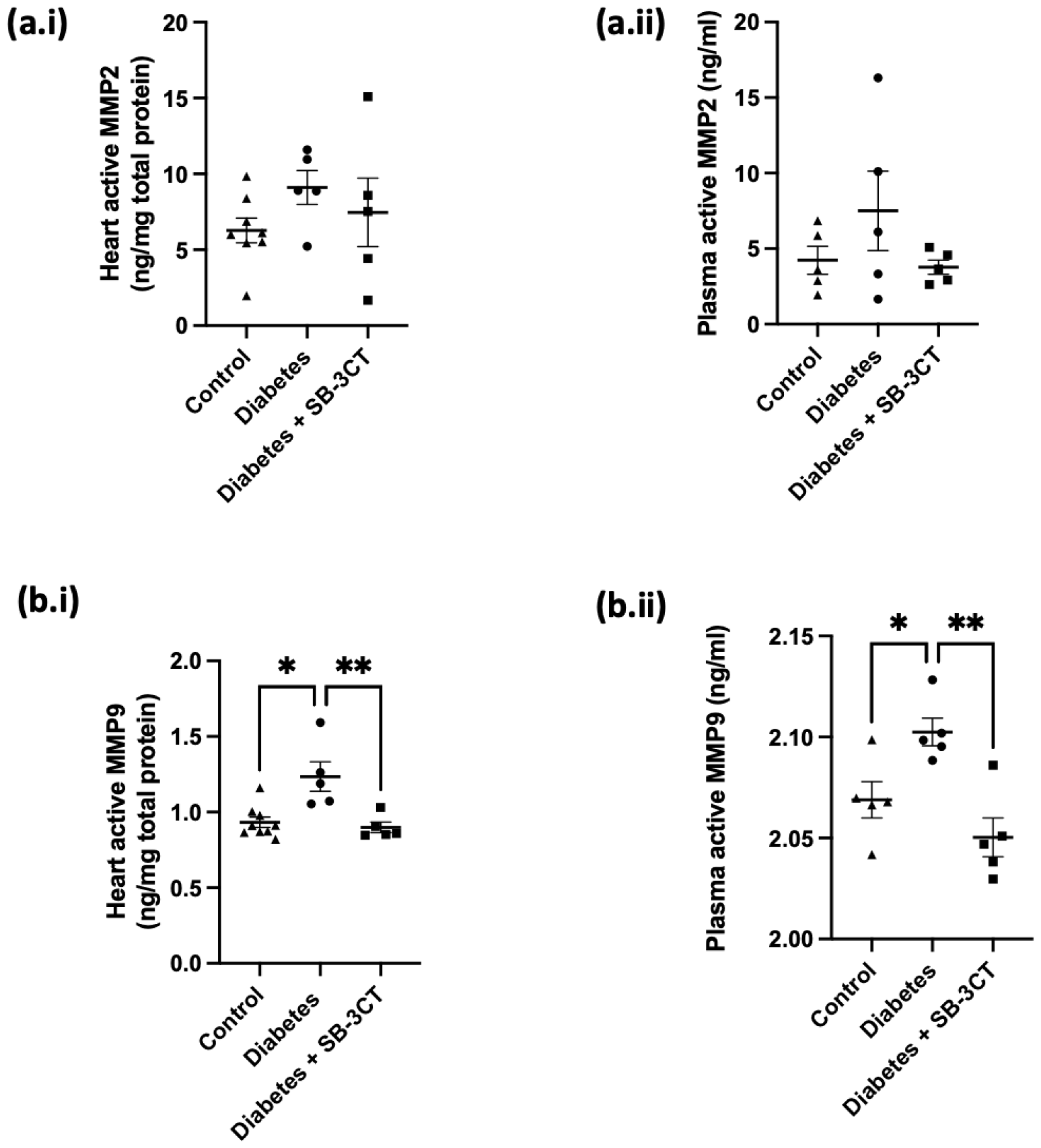
MMP9 activity is increased in diabetes and reduced with SB-3CT. Nine weeks post the induction of diabetes, plasma and mouse hearts were collected to assess MMP9 activity. (a.i) MMP2 activity in mouse heart tissue 9 weeks post diabetes induction (n=8 for control, and 5 for diabetes and diabetes + SB-3CT; control vs diabetes, p=0.39; diabetes vs diabetes + SB-3CT, p=0.69). (a.ii) MMP2 activity in mouse plasma (n=5 for all groups; control vs diabetes, p=0.36; diabetes vs diabetes + SB-3CT, p=0.28). (bi) MMP9 activity in mouse heart tissue 9 weeks post diabetes induction (n=9 for control, and 5 for diabetes and diabetes + SB-3CT; control vs diabetes, *p<0.05; diabetes vs diabetes + SB-3CT, **p<0.01; Kruskal-Wallis test). (b.ii) MMP9 activity in mouse plasma 9 weeks post diabetes induction (n=5 for all groups; control vs diabetes, *p<0.05; diabetes vs diabetes + SB-3CT, **p<0.01; One-way ANOVA)

### MMP9 activity is increased in diabetes and reduced with SB-3CT

To confirm that MMP activity was increased in our mouse model, the activity of MMP2 and 9 was assessed in both the plasma and heart. No significant difference in MMP2 activity was found in the heart and plasma of the mice which supports our previous findings [30]. As expected, a significant increase in MMP9 activity was found in the heart tissue and plasma of diabetic mice 9 weeks post the induction of diabetes. This was reduced when treated with SB-3CT proving the inhibitor to be effective.

### MMP2/9 inhibition prevents SDC4 shedding from ciCMVEC

To extend the relevance of the work to humans, we used human cells to investigate the mechanism of EGlx shedding. SDC4 is a common core protein of the EGlx and a known substrate for MMP9 [31]. A relationship has previously been established between TNF-α (a common proinflammatory cytokine upregulated in diabetes) and SDC4 shedding in glomerular endothelial cells [31]. Therefore, this was investigated using ciCMVEC. TNF-α caused a loss of cell surface-associated SDC4 compared to controls, when normalised to DAPI (Fig. 6a.ii). TNF-α also resulted in an increase in SDC4 mRNA expression (Fig. 6b) which may be a compensatory mechanism as a result of its shedding. Treatment with SB-3CT ameliorated these changes. A SDC4 ELISA (Fig. 6c) and Alcian blue assay (Fig. 6d) was conducted on the conditioned media to quantify SDC4 and GAG concentration as an indication of EGlx shedding. TNF-α resulted in an increased SDC4 and GAG concentration in the conditioned media compared to controls and this was significantly reduced by the MMP inhibitor.

**Fig. 6.**
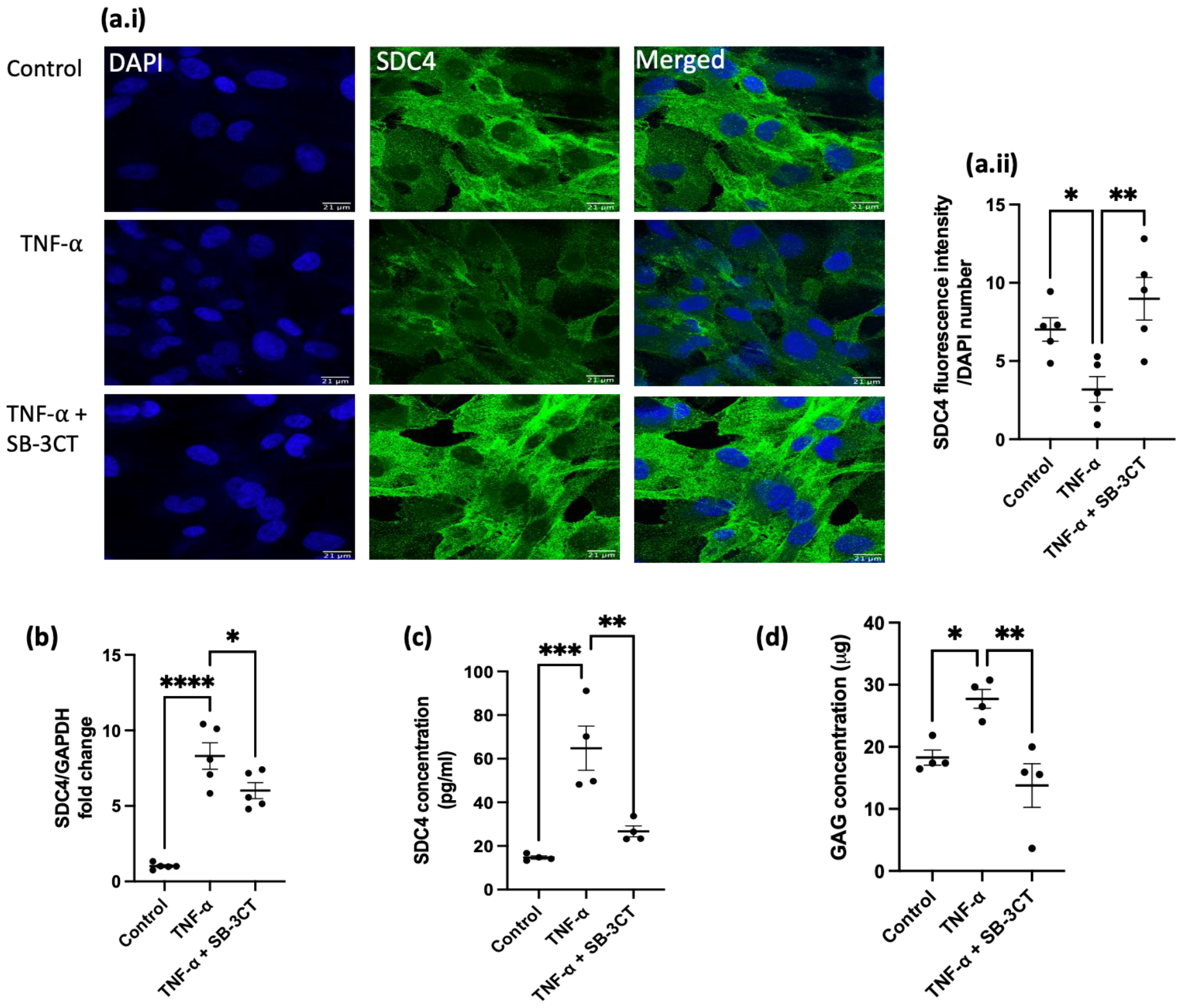
Inhibition of MMPs prevents SDC4 shedding from CMVEC. Cells were pre-incubated with SB-3CT for 2 hours before being treated with TNF-α for 6 hours. (a) SDC4 cell surface expression. Fluorescence intensity was normalized to cell number (n=5 for all groups; control vs TNF-α, *p<0.05; TNF-α vs TNF-α + SB-3CT, **p<0.01; One-way ANOVA). (b) SDC4 mRNA expression (n=5 for all groups; Control vs TNF-α, ****p<0.0001; TNF-α vs TNF-α + SB-3CT, *P<0.05; One-way ANOVA). (c) SDC4 concentration in the conditioned media (n=4 for all groups; Control vs TNF-α, ***p<0.001; TNF-α vs TNF-α + SB-3CT, **p<0.01; One-way ANOVA). (d) GAG concentration in the conditioned media (n=4 for all groups; Control vs TNF-α, *p<0.05; TNF-α vs TNF-α + SB-3CT, **p<0.01; One-way ANOVA)

### MMP9 causes SDC4 shedding from coronary microvascular endothelial cells

The direct role of MMP9 on the SDC4 shedding from ciCMVEC was investigated. As expected, TNF-α upregulated MMP9 mRNA expression (Fig. 7a) as well as resulted in an increase in MMP9 activity in the conditioned media (Fig. 7b). These were both reduced with SB-3CT. Treatment of ciCMVEC with MMP9 shRNA led to a reduction in MMP9 mRNA expression (Fig. 7c) confirming a successful knockdown. Knockdown of MMP9 expression significantly reduced SDC4 mRNA upregulation by TNF-α (Fig. 7d) and SDC4 concentration in the conditioned media suggesting reduced shedding (Fig. 7e).

**Fig. 7.**
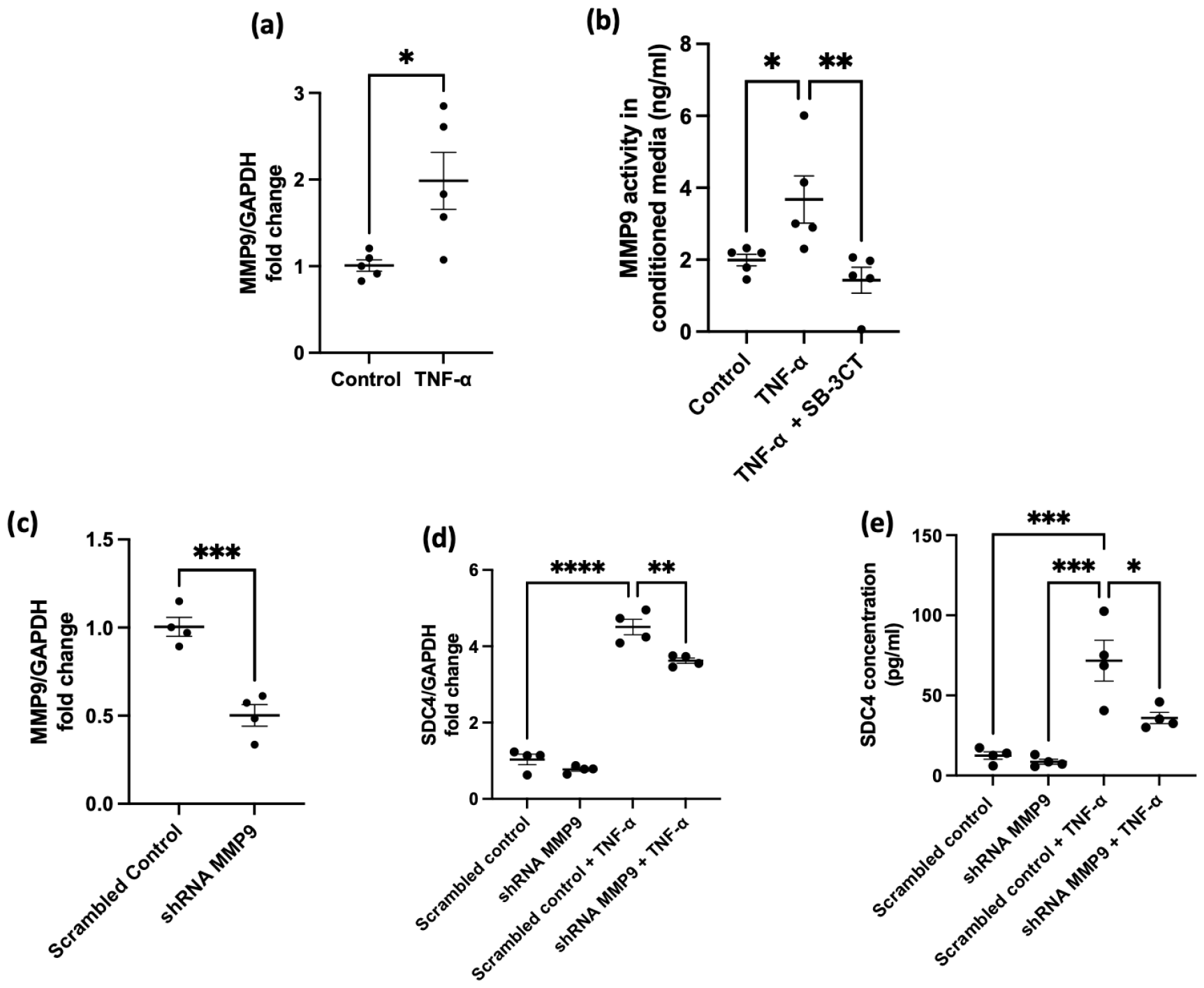
MMP9 causes SDC4 shedding from coronary microvascular endothelial cells. (a) MMP9 mRNA expression (n=5 for both groups; *p<0.05; unpaired t test). (b) MMP9 activity in the conditioned media (n=5 for all groups; control vs TNF-α, *p<0.05; TNF-α vs TNF-α + SB-3CT, **p< 0.01; One-way ANOVA). (c) MMP9 mRNA expression from shRNA MMP9 cells (n=4 for both groups; **p<0.01; unpaired t test). (d) SDC4 mRNA expression (n=4 for all groups; scrambled control vs scrambled control + TNF-α, ****p<0.0001. Scrambled control + TNF-α vs shRNA MMP9 + TNF-α, **p<0.01; One-way ANOVA). No significant difference found between shRNA MMP9 and scrambled control (p=0.5). (e) SDC4 concentration in the conditioned media (n=4 for all groups; scrambled control vs scrambled control + TNF-α, ***p<0.001. Scrambled control + TNF-α vs shRNA MMP9 + TNF-α, *p<0.05)

## Discussion

We have shown that MMP9 causes shedding of the coronary microvascular EGlx and that inhibition of MMP2/9 protects the EGlx and restores diastolic function in a mouse model of DCM. The EGlx is a vital regulator of permeability and we have previously shown that removal of the EGlx from human CMVEC in vitro results in increased albumin permeability [30]. We provide further support to this through the observation of increased albumin in the heart tissue of diabetic mice when compared to the controls. Research suggests that an increased permeability can directly cause diastolic dysfunction [1] and therefore the EGlx is an important target of protection for the heart. The reduction in EGlx depth observed in this study is likely to result in increased microvascular permeability leading to a rise in interstitial pressures and stiffness of the myocardium [1, 9].

In this model, diastolic dysfunction occurred in the absence of systolic function, and this was improved with MMP2/9 inhibition. In the early phase of DCM (subclinical period), patients present with impaired diastolic function (indicated by a reduced E/A ratio) [21, 43] and normal systolic function. As this progresses, patients will eventually develop heart failure with limited ejection fraction [44]. Echocardiography is widely used clinically to assess diastolic function. Observing diastolic dysfunction before treatment with SB-3CT allows us to identify the inhibition of MMPs as a treatment rather than prevention of DCM. This provides more translatability to this study as clinically patients are diagnosed with DCM before being treated and so this study has therapeutic value. At 9 weeks post the induction of diabetes, no systolic dysfunction was observed but a reduced E/A ratio along with an increased IVRT was found suggesting diastolic dysfunction. This result is in line with previous studies, which at 10-12 weeks post STZ, do not recognise any changes in systolic function but an increase in IVRT and reduced E/A ratio in diabetes [12, 30, 44].

Methods for investigating the EGlx have greatly evolved. We have previously validated the use of lectin labelling for EGlx measurements against the gold standard electron microscopy measurements in both the heart and the kidneys [5, 30]. Using lectin staining in conjunction with peak-to-peak analysis, we have shown a reduction in EGlx depth in the diabetic group when compared to the control, consistent with previous results [28–30, 32]. Treatment with SB-3CT was able to protect the EGlx of the heart and therefore, we have successfully identified MMPs as a therapeutic target of inhibition to protect the coronary microvascular EGlx. The use of lectins allows the direct visualisation of the EGlx and is increasingly being used in EGlx research by binding to disaccharide moieties on the GAG side. It is known that GAGs are distributed in areas other than the EGlx such as the basement membrane, and connective tissues [24]. We overcome this with the utilisation of the peak-to-peak method of assessing EGlx depth. As the EGlx is on the luminal side, only the fluorescent peak from the lectin that occurs before the membrane is taken as the EGlx. The mechanism highlighted in this study is that an increase in MMP9 activity causes the excessive shedding of the coronary microvascular EGlx in DCM. This results in an increase in microvascular permeability, leading to left ventricular stiffness and diastolic dysfunction. We show that MMP9 activity is upregulated in both the heart tissue and plasma. Inhibition of MMP9 improved EGlx depth and restored diastolic function in diabetic mice. The importance of MMP9 on EGlx shedding is further demonstrated by MMP9 knockdown in ciCMVEC. The significant reduction in SDC4 concentration in the conditioned media of cells with reduced MMP9 expression, when treated with TNF-α, shows that MMP9 directly contributes to the shedding of SDC4 from CMVEC.

DCM is complex with various changes occurring to the physiological state, including an increase in proinflammatory cytokines [39]. Indeed, it is recognised that hyperglycaemia induces proinflammatory cytokines such as TNF-α, IL-1b, and IL-6 all of which contribute to the complex mechanisms of diabetes [16, 37].

Focusing on a specific component of the pathophysiology of diabetes has allowed us to better understand and elucidate the mechanisms of EGlx damage in diabetes. In this study, we focused on TNF-α and identified that TNF-α causes the MMP9-mediated shedding of SDC4. We have recently shown that SDC4 is reduced at coronary microvessels as a result of DCM [30]. Therefore, this study not only supports this but identifies a mechanism that may lead to reduced endothelial SDC4 in the heart during DCM.

The findings of our study hold significant clinical importance in the context of DCM. Identifying MMPs as a shedder of the EGlx in DCM opens new avenues for therapeutic interventions to improve patient outcomes. In this study, we have reinforced that the EGlx is damaged in DCM and that its protection is of paramount importance. By inhibiting MMP2/9 with SB-3CT, we observed a restoration of diastolic function, protection of EGlx depth and a reduction in albumin permeability. Patient plasma MMP9 activity levels and EGlx integrity could serve as potential biomarkers to assess disease severity. Integrating these assessments into clinical practice may facilitate early detection and personalised management of DCM.

In this study we did not observe a significant difference in MMP2 between the groups and thus emphasis is placed more on MMP9 in which the activity was evidently upregulated in diabetes and reduced with SB-3CT. It would be beneficial to confidently identify a specific MMP that can be targeted to protect the EGlx in diabetes to enhance the development of therapeutics and reduce the side effects observed from broad spectrum MMP inhibitors. In this study we have started to identify MMP9 as a key target to protect the coronary microvascular EGlx. Therefore, future studies will aim to further investigate this specific MMP to increase the potential of targeted therapeutics.

We acknowledge the limitation of the study in that only a type 1 model of diabetes was utilised. However, we have now provided the groundwork for future studies in which this work can applied to a type 2 model of diabetes. As we have previously shown that protecting the coronary microvascular EGlx is beneficial in both type 1 and type 2 models of DCM [30], there is confidence that this work can successfully be applied to type 2 models and thus warrants investigation. In fact, an increase in MMP9 activity has previously been recorded in type 2 diabetic patients and animal models [4, 10, 25] providing support to the idea that MMP inhibition may protect the EGlx in type 2 diabetes similarly to what we observe in our type 1 model.

Our study provides valuable insights into the potential therapeutic implications of MMP inhibition in protecting the EGlx and improving cardiac function in DCM. Currently, treatments for DCM are limited, and there is a critical need for novel therapeutic approaches. The use of SB-3CT to inhibit MMP9 has proven successful in many preclinical studies [11, 14, 17, 18, 40, 45] and now we have shown it to be effective in inhibiting MMP9 in a mouse model of DCM. Our study provides evidence that specific MMP9 inhibition could be a promising strategy to protect the EGlx and restore cardiac function. Developing targeted therapies to preserve the EGlx in diabetes could revolutionize the management of DCM and improve patient outcomes.

In conclusion, our study underscores the clinical importance of MMP9 in the context of DCM and its impact on the EGlx. The findings support the rationale for targeting MMP9 to protect the EGlx and improve diastolic function. By elucidating this specific mechanism, our research contributes to the development of targeted therapeutic approaches for DCM and highlights the preservation of the EGlx as a valuable strategy in managing cardiovascular complications in diabetes.

## Data Availability

The data underlying this article are available in the article.

## Acknowledgements

The authors gratefully acknowledge the Wolfson Bioimaging facility (University of Bristol, UK) for their support and assistance in this work.

This work was funded by a British Heart Foundation (BHF) Integrative Cardiovascular Science PhD programme studentship (FS/19/53/34887A) and a BHF project grant (PG/20/7/34849).

## Notes

### Competing Interest Statement

The authors have declared no competing interest.

## References

1. Abelanet A, Camoin M, Rubin S, Bougaran P, Delobel V, Pernot M, Forfar I, Guilbeau-Frugier C, Galès C, Bats ML, Renault M-A, Dufourcq P, Couffinhal T, Duplàa C (2022) Increased Capillary Permeability in Heart Induces Diastolic Dysfunction Independently of Inflammation, Fibrosis, or Cardiomyocyte Dysfunction. Arterioscler Thromb Vasc Biol 42:745–763. doi: 10.1161/ATVBAHA.121.317319

2. Betteridge KB, Arkill KP, Neal CR, Harper SJ, Foster RR, Satchell SC, Bates DO, Salmon AHJ (2017) Sialic acids regulate microvessel permeability, revealed by novel in vivo studies of endothelial glycocalyx structure and function. J Physiol 595:5015–5035. doi: 10.1113/JP274167

3. Cai H, Ma Y, Jiang L, Mu Z, Jiang Z, Chen X, Wang Y, Yang G-Y, Zhang Z (2017) Hypoxia Response Element-Regulated MMP-9 Promotes Neurological Recovery via Glial Scar Degradation and Angiogenesis in Delayed Stroke. Mol Ther 25:1448–1459. doi: 10.1016/j.ymthe.2017.03.020

4. Collin HL, Sorsa T, Meurman JH, Niskanen L, Salo T, Rönkä H, Konttinen YT, Koivisto AM, Uusitupa M (2000) Salivary matrix metalloproteinase (MMP-8) levels and gelatinase (MMP-9) activities in patients with type 2 diabetes mellitus. J Periodontal Res 35:259–265. doi: 10.1034/j.1600-0765.2000.035005259.x

5. Crompton M, Ferguson JK, Ramnath R, Onions KL, Ogier AS, Gamez M, Down CJ, Skinner LJ, Wong KHF, Dixon LK, Sutak J, Harper SJ, Pontrelli P, Gesualdo L, Heerspink HL, Toto RD, Welsh GI, Foster RR, Satchell SC, Butler MJ (2023) Mineralocorticoid receptor antagonism in diabetes reduces albuminuria by preserving the glomerular endothelial glycocalyx. JCI Insight 8:154164–154180. doi: 10.1172/jci.insight.154164

6. De Blasio MJ, Huynh N, Deo M, Dubrana LE, Walsh J, Willis A, Prakoso D, Kiriazis H, Donner DG, Chatham JC, Ritchie RH (2020) Defining the Progression of Diabetic Cardiomyopathy in a Mouse Model of Type 1 Diabetes. Front Physiol 11:124–139. doi: 10.3389/fphys.2020.00124

7. Dede E, Liapis D, Davos C, Katsimpoulas M, Varela A, Mpotis I, Kostomitsopoulos N, Kadoglou NPE (2022) The effects of exercise training on cardiac matrix metalloproteinases activity and cardiac function in mice with diabetic cardiomyopathy. Biochem Biophys Res Commun 586:8–13. doi: 10.1016/j.bbrc.2021.11.013

8. Dia M, Gomez L, Thibault H, Tessier N, Leon C, Chouabe C, Ducreux S, Gallo-Bona N, Tubbs E, Bendridi N, Chanon S, Leray A, Belmudes L, Couté Y, Kurdi M, Ovize M, Rieusset J, Paillard M (2020) Reduced reticulum–mitochondria Ca2+ transfer is an early and reversible trigger of mitochondrial dysfunctions in diabetic cardiomyopathy. Basic Res Cardiol 115:74–94. doi: 10.1007/s00395-020-00835-7

9. Dongaonkar RM, Stewart RH, Geissler HJ, Laine GA (2010) Myocardial microvascular permeability, interstitial oedema, and compromised cardiac function. Cardiovasc Res 87:331–339. doi: 10.1093/cvr/cvq145

10. Gonçalves FM, Jacob-Ferreira ALB, Gomes VA, Casella-Filho A, Chagas ACP, Marcaccini AM, Gerlach RF, Tanus-Santos JE (2009) Increased circulating levels of matrix metalloproteinase (MMP)-8, MMP-9, and pro-inflammatory markers in patients with metabolic syndrome. Clin Chim Acta 403:173–177. doi: 10.1016/j.cca.2009.02.013

11. Gooyit M, Suckow MA, Schroeder VA, Wolter WR, Mobashery S, Chang M (2012) Selective Gelatinase Inhibitor Neuroprotective Agents Cross the Blood-Brain Barrier. ACS Chem Neurosci 3:730–736. doi: 10.1021/cn300062w

12. Hou N, Mai Y, Qiu X, Yuan W, Li Y, Luo C, Liu Y, Zhang G, Zhao G, Luo J (2019) Carvacrol Attenuates Diabetic Cardiomyopathy by Modulating the PI3K/AKT/GLUT4 Pathway in Diabetic Mice. Front Pharmacol 10:998–1012

13. Jang H (2018) Regulation of Cyclic AMP-Response Element Binding Protein Zhangfei (CREBZF) Expression by Estrogen in Mouse Uterus. Dev Reprod 22:95–104. doi: 10.12717/DR.2018.22.1.095

14. Jia F, Yin YH, Gao GY, Wang Y, Cen L, Jiang J (2014) MMP-9 Inhibitor SB-3CT Attenuates Behavioral Impairments and Hippocampal Loss after Traumatic Brain Injury in Rat. J Neurotrauma 31:1225–1234. doi: 10.1089/neu.2013.3230

15. Kannel WB, Hjortland M, Castelli WP (1974) Role of diabetes in congestive heart failure: The Framingham study. Am J Cardiol 34:29–34. doi: 10.1016/0002-9149(74)90089-7

16. Kaur N, Guan Y, Raja R, Ruiz-Velasco A, Liu W (2021) Mechanisms and Therapeutic Prospects of Diabetic Cardiomyopathy Through the Inflammatory Response. Front Physiol 12:694864–694874. doi: 10.3389/fphys.2021.694864

17. Kleifeld O, Kotra LP, Gervasi DC, Brown S, Bernardo MM, Fridman R, Mobashery S, Sagi I (2001) X-ray Absorption Studies of Human Matrix Metalloproteinase-2 (MMP-2) Bound to a Highly Selective Mechanism-based Inhibitor: COMPARISON WITH THE LATENT AND ACTIVE FORMS OF THE ENZYME *. J Biol Chem 276:17125–17131. doi: 10.1074/jbc.M011604200

18. Lee M, Chen Z, Tomlinson BN, Gooyit M, Hesek D, Juárez MR, Nizam R, Boggess B, Lastochkin E, Schroeder VA, Wolter WR, Suckow MA, Cui J, Mobashery S, Gu Z, Chang M (2015) Water-Soluble MMP-9 Inhibitor Reduces Lesion Volume after Severe Traumatic Brain Injury. ACS Chem Neurosci 6:1658–1664. doi: 10.1021/acschemneuro.5b00140

19. Li W, Wang W (2018) Structural alteration of the endothelial glycocalyx: contribution of the actin cytoskeleton. Biomech Model Mechanobiol 17:147–158. doi: 10.1007/s10237-017-0950-2

20. Liew H, Roberts MA, Pope A, McMahon LP (2021) Endothelial glycocalyx damage in kidney disease correlates with uraemic toxins and endothelial dysfunction. BMC Nephrol 22:21–29. doi: 10.1186/s12882-020-02219-4

21. Liu JE, Palmieri V, Roman MJ, Bella JN, Fabsitz R, Howard BV, Welty TK, Lee ET, Devereux RB (2001) The impact of diabetes on left ventricular filling pattern in normotensive and hypertensive adults: the strong heart study. J Am Coll Cardiol 37:1943–1949. doi: 10.1016/S0735-1097(01)01230-X

22. Liu X, Xu Q, Wang X, Zhao Z, Zhang L, Zhong L, Li L, Kang W, Zhang Y, Ge Z (2015) Irbesartan ameliorates diabetic cardiomyopathy by regulating protein kinase D and ER stress activation in a type 2 diabetes rat model. Pharmacol Res 93:43–51. doi: 10.1016/j.phrs.2015.01.001

23. Lorenzo-Almorós A, Tuñón J, Orejas M, Cortés M, Egido J, Lorenzo Ó (2017) Diagnostic approaches for diabetic cardiomyopathy. Cardiovasc Diabetol 16:28. doi: 10.1186/s12933-017-0506-x

24. Lubec G (1982) [Glycosaminoglycan-glomerular basement membrane interactions revealed by affinity chromatography (author’s transl)]. Padiatr Padol 17:591–596

25. Luo S, Li W, Wu W, Shi Q (2021) Elevated expression of MMP8 and MMP9 contributes to diabetic osteoarthritis progression in a rat model. J Orthop Surg 16:64. doi: 10.1186/s13018-021-02208-9

26. Mierzejewski M, Paplinska-Goryca M, Korczynski P, Krenke R (2021) Primary human mesothelial cell culture in the evaluation of the inflammatory response to different sclerosing agents used for pleurodesis. Physiol Rep 9:e14846–e14855. doi: 10.14814/phy2.14846

27. Moore KH, Murphy HA, George EM (2021) The glycocalyx: a central regulator of vascular function. Am J Physiol-Regul Integr Comp Physiol 320:R508–R518. doi: 10.1152/ajpregu.00340.2020

28. Nieuwdorp M, Haeften TW van, Gouverneur MCLG, Mooij HL, Lieshout MHP van, Levi M, Meijers JCM, Holleman F, Hoekstra JBL, Vink H, Kastelein JJP, Stroes ESG (2006) Loss of Endothelial Glycocalyx During Acute Hyperglycemia Coincides With Endothelial Dysfunction and Coagulation Activation In Vivo. Diabetes 55:480–486. doi: 10.2337/diabetes.55.02.06.db05-1103

29. Nieuwdorp M, Mooij HL, Kroon J, Atasever B, Spaan JAE, Ince C, Holleman F, Diamant M, Heine RJ, Hoekstra JBL, Kastelein JJP, Stroes ESG, Vink H (2006) Endothelial Glycocalyx Damage Coincides With Microalbuminuria in Type 1 Diabetes. Diabetes 55:1127–1132. doi: 10.2337/diabetes.55.04.06.db05-1619

30. Qiu Y, Buffonge S, Ramnath R, Jenner S, Fawaz S, Arkill KP, Neal C, Verkade P, White SJ, Hezzell M, Salmon AHJ, Suleiman M-S, Welsh GI, Foster RR, Madeddu P, Satchell SC (2022) Endothelial glycocalyx is damaged in diabetic cardiomyopathy: angiopoietin 1 restores glycocalyx and improves diastolic function in mice. Diabetologia 65:879–894. doi: 10.1007/s00125-022-05650-4

31. Ramnath R, Foster RR, Qiu Y, Cope G, Butler MJ, Salmon AH, Mathieson PW, Coward RJ, Welsh GI, Satchell SC (2014) Matrix metalloproteinase 9-mediated shedding of syndecan 4 in response to tumor necrosis factor α: a contributor to endothelial cell glycocalyx dysfunction. FASEB J 28:4686–4699. doi: 10.1096/fj.14-252221

32. Ramnath RD, Butler MJ, Newman G, Desideri S, Russell A, Lay AC, Neal CR, Qiu Y, Fawaz S, Onions KL, Gamez M, Crompton M, Michie C, Finch N, Coward RJ, Welsh GI, Foster RR, Satchell SC (2020) Blocking matrix metalloproteinase-mediated syndecan-4 shedding restores the endothelial glycocalyx and glomerular filtration barrier function in early diabetic kidney disease. Kidney Int 97:951–965. doi: 10.1016/j.kint.2019.09.035

33. Reitsma S, Slaaf DW, Vink H, van Zandvoort MAMJ, oude Egbrink MGA (2007) The endothelial glycocalyx: composition, functions, and visualization. Pflugers Arch 454:345–359. doi: 10.1007/s00424-007-0212-8

34. Rubler S, Dlugash J, Yuceoglu YZ, Kumral T, Branwood AW, Grishman A (1972) New type of cardiomyopathy associated with diabetic glomerulosclerosis. Am J Cardiol 30:595–602. doi: 10.1016/0002-9149(72)90595-4

35. Satchell SC, Tasman CH, Singh A, Ni L, Geelen J, von Ruhland CJ, O’Hare MJ, Saleem MA, van den Heuvel LP, Mathieson PW (2006) Conditionally immortalized human glomerular endothelial cells expressing fenestrations in response to VEGF. Kidney Int 69:1633–1640. doi: 10.1038/sj.ki.5000277

36. Schindelin J, Arganda-Carreras I, Frise E, Kaynig V, Longair M, Pietzsch T, Preibisch S, Rueden C, Saalfeld S, Schmid B, Tinevez J-Y, White DJ, Hartenstein V, Eliceiri K, Tomancak P, Cardona A (2012) Fiji: an open-source platform for biological-image analysis. Nat Methods 9:676–682. doi: 10.1038/nmeth.2019

37. Stentz FB, Umpierrez GE, Cuervo R, Kitabchi AE (2004) Proinflammatory Cytokines, Markers of Cardiovascular Risks, Oxidative Stress, and Lipid Peroxidation in Patients With Hyperglycemic Crises. Diabetes 53:2079–2086. doi: 10.2337/diabetes.53.8.2079

38. Weng J, Trinh S, Lee R, Metwale R, Sharma A (2022) Impact of High Glucose on Ocular Surface Glycocalyx Components: Implications for Diabetes-Associated Ocular Surface Damage. Int J Mol Sci 23:14289–14301. doi: 10.3390/ijms232214289

39. Westermann D, Van Linthout S, Dhayat S, Dhayat N, Schmidt A, Noutsias M, Song X-Y, Spillmann F, Riad A, Schultheiss H-P, Tschöpe C (2007) Tumor necrosis factor-alpha antagonism protects from myocardial inflammation and fibrosis in experimental diabetic cardiomyopathy. Basic Res Cardiol 102:500–507. doi: 10.1007/s00395-007-0673-0

40. Wu M-Y, Gao F, Yang X-M, Qin X, Chen G-Z, Li D, Dang B-Q, Chen G (2020) Matrix metalloproteinase-9 regulates the blood brain barrier via the hedgehog pathway in a rat model of traumatic brain injury. Brain Res 1727:146553–146563. doi: 10.1016/j.brainres.2019.146553

41. Ye Y, Kuang X, Xie Z, Liang L, Zhang Z, Zhang Y, Ma F, Gao Q, Chang R, Lee H-H, Zhao S, Su J, Li H, Peng J, Chen H, Yin M, Peng C, Yang N, Wang J, Liu J, Liu H, Han L, Chen X (2020) Small-molecule MMP2/MMP9 inhibitor SB-3CT modulates tumor immune surveillance by regulating PD-L1. Genome Med 12:83–101. doi: 10.1186/s13073-020-00780-z

42. Yilmaz O, Afsar B, Ortiz A, Kanbay M (2019) The role of endothelial glycocalyx in health and disease. Clin Kidney J 12:611–619. doi: 10.1093/ckj/sfz042

43. Zarich SW, Arbuckle BE, Cohen LR, Roberts M, Nesto RW (1988) Diastolic abnormalities in young asymptomatic diabetic patients assessed by pulsed Doppler echocardiography. J Am Coll Cardiol 12:114–120. doi: 10.1016/0735-1097(88)90364-6

44. Zhang X, Ke P-X, Yuan X, Zhang G-P, Chen W-L, Zhang G-S (2021) Forskolin Protected against Streptozotocin-Induced Diabetic Cardiomyopathy via Inhibition of Oxidative Stress and Cardiac Fibrosis in Mice. BioMed Res Int 2021:8881843–8881850. doi: 10.1155/2021/8881843

45. Zhou P, Yang C, Zhang S, Ke Z-X, Chen D-X, Li Y-Q, Li Q (2021) The Imbalance of MMP-2/TIMP-2 and MMP-9/TIMP-1 Contributes to Collagen Deposition Disorder in Diabetic Non-Injured Skin. Front Endocrinol 12:734485–734495. doi: 10.3389/fendo.2021.734485

46. (2013) Selective Inhibition of Matrix Metalloproteinase-9 Attenuates Secondary Damage Resulting from Severe Traumatic Brain Injury. PLOS ONE 8:e76904–e16917. doi: 10.1371/journal.pone.0076904

